# Multi-modal Phantom Experiments, mimicking Flow through the Mitral Heart Valve

**DOI:** 10.1101/2024.02.20.581131

**Authors:** Lea Christierson, Petter Frieberg, Tania Lala, Johannes Töger, Petru Liuba, Johan Revstedt, Hanna Isaksson, Nina Hakacova

## Abstract

**Purpose:** Fluid-structure interaction (FSI) models are more commonly applied in medical research as computational power is increasing. However, understanding the accuracy of FSI models is crucial, especially in the context of heart valve disease in patient-specific models. Therefore, this study aimed to create a multi-modal benchmarking data set for FSI models, based on clinically important parameters, such as the pressure, velocity, and valve opening, with an *in vitro* phantom setup.

**Method:** An *in vitro* setup was developed with a 3D-printed phantom mimicking the left heart, including a deforming mitral valve. A range of pulsatile flows was created with a computer-controlled motor-and-pump setup. Invasive catheter measurements, magnetic resonance imaging (MRI), and echocardiography (Echo) imaging were used to measure pressure and velocity in the domain. Furthermore, the valve opening was quantified based on cine MRI and Echo images.

**Result:** The experimental setup, with 0.5 % cycle-to-cycle variation, was successfully built and six different flow cases were investigated. Higher velocity through the mitral valve was observed for increased cardiac output. The pressure difference across the valve also followed this trend. The flow in the phantom was qualitatively assessed by the velocity profile in the ventricle and by streamlines obtained from 4D phase-contrast MRI.

**Conclusion:** A multi-modal set of validation data for FSI models has been created, based on parameters relevant for diagnosis of heart valve disease. All data is publicly available for future development of computational heart valve models.

## INTRODUCTION

In recent years, fluid-structure interaction (FSI) simulations have become increasingly prevalent as computational resources have advanced, particularly in the field of medical research and heart valve disease [1]. Assessing the accuracy and reliability of FSI simulations is a complex task since it models both the fluid and structural domains [2], particularly in a clinical context where it is essential to determine the extent of their impact on diagnosis. Understanding the degree of accuracy of FSI simulations in a clinical setting is important to ensure reliable results that support medical decision-making. To establish the accuracy of simulations, validation against experimental data is a fundamental step to ensure reliable results, usable for clinicians [3].

Patient-specific FSI simulations of heart valve disease can enable a better understanding of the valvular dynamics and complement traditional imaging modalities [1]. Furthermore, they have the potential to predict the valve repair outcome and enhance personalized treatment, since a digital model can help clinicians improve planning surgical or transcatheter interventions [4]. Despite their potential benefits, these simulation models have not yet been implemented in clinical practice, partly due to the lack of comprehensive validation [5].

Previous FSI simulation studies on heart valves have compared their simulation results to previous publications [6]–[9], to assess their agreement with other numeric data. While this approach is a common form of evaluation, it has limitations since the model accuracy is not measured against experimental data [3]. Only a few studies have developed a setup for experimental validation of FSI simulations [3], [10], [11]. Most commonly, parameters relevant to the development of numerical methods have been investigated [12]–[17], such as tissue strain and stress, vortex formation, and leaflet angular velocity. However, there is a lack of investigation into parameters relevant to diagnosis of heart valve disease, particularly the pressure gradient and the valve opening orifice dimensions.

Therefore, this study aimed to create benchmarking data for FSI simulation models, on an *in vitro* setup mimicking the left heart including a deforming mitral valve. A comprehensive multi-modal validation, with invasive pressure measurements, magnetic resonance imaging (MRI), and echocardiographic (Echo) imaging, was employed. This study aimed to measure and quantify key parameters such as ventricular and atrial pressure, velocity within the domain, and mitral valve opening to provide extensive validation data for future development of FSI models. The data is provided openly for use by the research community [18].

## MATERIALS AND METHODS

In this study, an *in vitro* test setup is presented (Fig. 1), where a phantom mimicking the left heart including the mitral valve, was used to create benchmarking data for the validation of computer-based models for future patient-specific simulations. Pulsatile physiological waveforms were produced with a computer-controlled motor and pump assembly. Invasive pressure measurements and medical imaging techniques were employed such that the pressure, velocity, and opening of the valve could be measured.

**Fig. 1.**
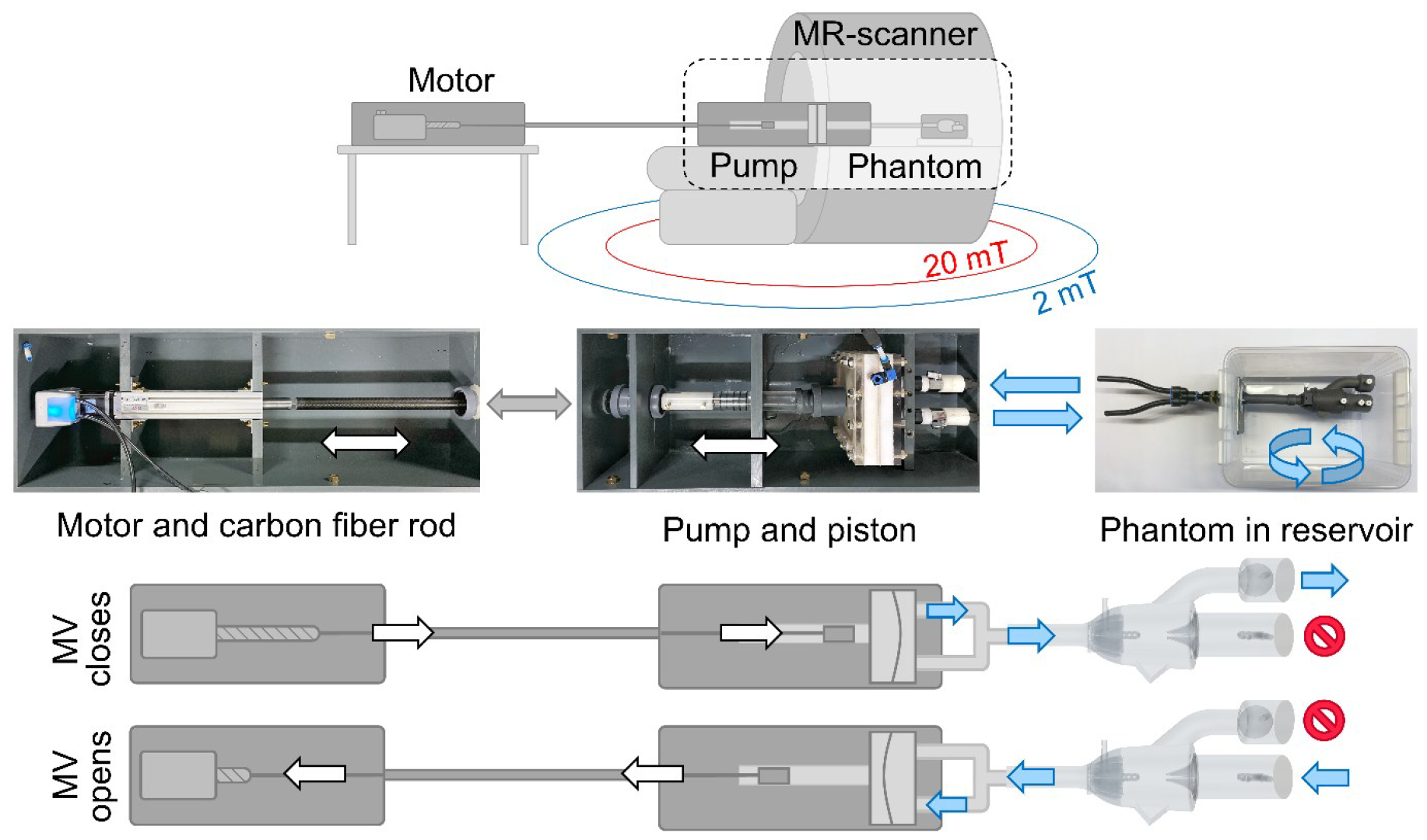
A schematic sketch of the pump setup within the MR scanner, with the 2 mT and 20 mT safety lines marked. The computer-controlled motor connected to a piston via a carbon fiber rod (grey arrow) is displayed along with sketches of the motor and pump configurations during the pump cycle causing the mitral valve (MV) to open and close. The piston pumps water in and out of the pump (white/black arrows), which in turn is connected to the phantom inside a plastic box, acting as a water reservoir. The blue arrows indicate the direction of the water.

### The in vitro test setup

The positive displacement pump [19] has one inlet and one outlet, that allowed water to be pumped out of a membrane chamber and back in, as the piston in the pump moved back and forth (Fig. 1). The setup has a stroke volume of 0-130 ml, both as a forward- and backward flow, with stroke rates of 1-200 beats per minute, thus a large variety of cardiac outputs (CO) can be imitated. The pump was programmed to generate three different waveforms with flows corresponding to a CO of 2.9, 4.4, and 5.4 l/min, at 60 beats per minute.

The pump was driven by a rod-style actuator with a 56B10A motor (C type) (Myostat Motion Control Inc., Newmarket, ON Canada). It was operated with Cool Muscle Language coding implemented in the software Control Room (Myostat Motion Control Inc., Newmarket, ON Canada). To transfer the force from the motor to the pump, a carbon fiber rod connection was used. The distance between the components of the pump setup was calculated to ensure a proper position inside the MR room; with the MR-safe pump located in the bore close to the phantom, at the iso-center, and with the ferromagnetic motor positioned outside the 2 mT safety line (Fig. 1). The Echo exam and pressure measurements were performed at a different time point, outside the MR room.

The phantom was connected to the pump at the apex and was mounted in a plastic box filled with 25 liters of room temperature water (Fig. 1), which acted as a reservoir that allowed water to freely enter and exit the model.

### Phantom geometry and flow description

A simplified *in vitro* model of the left heart was developed to ensure a controllable experimental setup and reliable measurements with few confounding factors. The phantom included a ventricle, atrium, mitral and aortic valves, and the apex (Fig. 2a-b). The mitral valve leaflets were designed to open and close without the aid of chordae tendineae or papillary muscles, as for the native mitral valve, but purely driven by the water flow. In five out of six cases, this worked well.

**Fig. 2.**
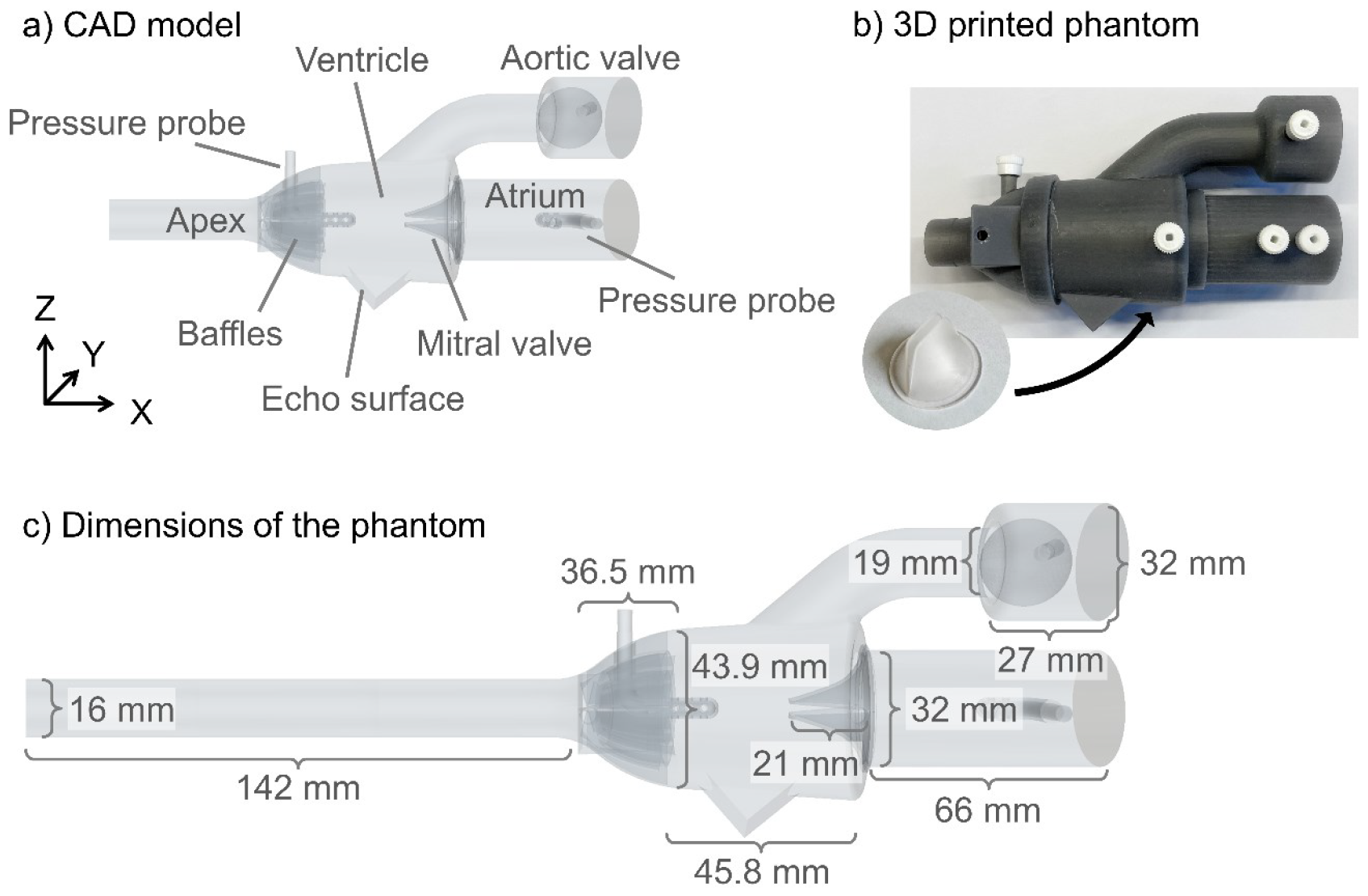
A schematic of the geometry of the phantom. a) The computer-aided design (CAD) geometry used for 3D printing, b) the 3D-printed phantom and a silicone mitral valve are shown. c) Some of the dimensions of the phantom and mitral valve. All exact measurements can be found in the CAD geometry files in the repository [18].

The flow entered and exited through the apex; when flow entered through the apex, the deformable mitral valve closed and the aortic valve, modeled as a ball valve, opened. As the flow was reversed and exited through the apex, the mitral valve opened, and the aortic ball valve closed. Thus, the flow in the beating heart was mimicked, and diastolic and systolic valve configurations were obtained despite the rigid phantom heart model (Fig. 1). A uniform waveform of the flow at the apex was maintained with baffles at the base of the heart to ensure an evenly distributed flow through the phantom (Fig. 2a).

Further, the phantom was equipped with flat surfaces for echo imaging, to ensure the same views were obtained for all investigated cases. The phantom also included two static pressure probes, reaching into the ventricle and atrium with a connection outside the model. Using Luer connections, catheters could be mounted to these for the pressure measurements (Fig. 2a). The dimensions of the phantom and the mitral valve approximate the dimensions of the human adult heart (Fig. 2c).

The phantom was 3D printed in a Form 2 printer (Formlabs Inc, Somerville, USA) with a stereolithography 3D printing technique in rigid (noncompliant) photopolymer resin (Grey Pro Resin, Formlabs Inc, Somerville, USA). The rigid flow domain was printed in such a way that the atrium was detachable from the ventricle, which enabled the silicone mitral valve to be changed. The heart valves were cast in a clear flexible silicone material (Agilus30, Stratasys Ltd., Eden Prairie, USA) with stiffness Shore A25, A40, and A60 (Dreve Biopor AB, Damvig A/S, Copenhagen, Denmark). Tensile tests on the silicone material were performed on dog bone samples for future material modeling purposes, reported in Appendix A.

### Measurements and analysis

#### Pressure catheters

The pressure in the atrium and ventricle was measured (Fig. 3) with fluid-filled catheters connected to a PowerLab Data Acquisition system (LabChart 7, ADInstruments, Dunedin, New Zealand). The catheters were connected to the pitot tube-like pressure probes of the phantom via Luer connections (Fig. 2) and the pressure at the atrium and ventricle was simultaneously recorded during 17 consecutive pump cycles. The pressure data was averaged over all cycles to reduce noise and cycle-to-cycle variations in the data.

**Fig. 3.**
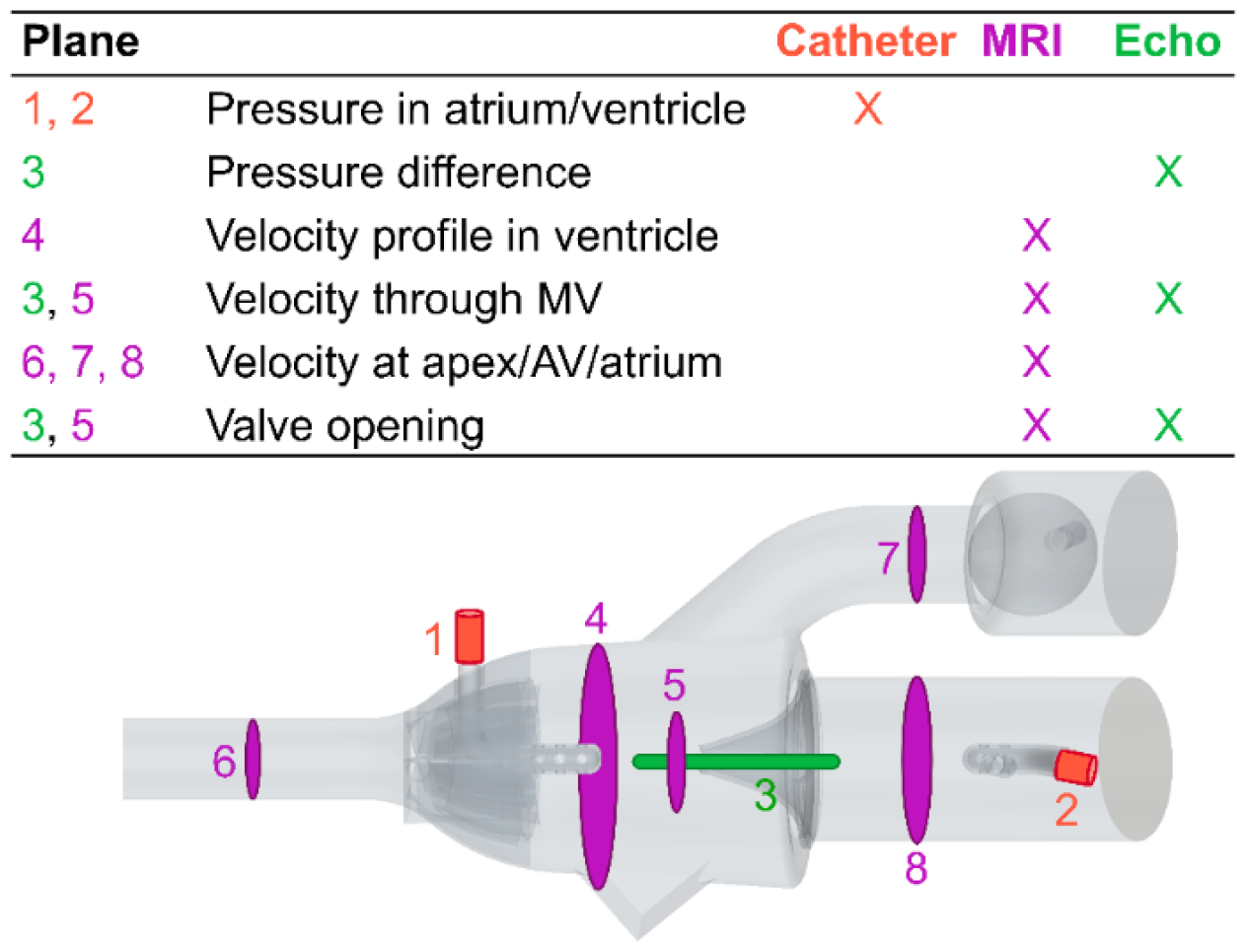
An overview of parameters measured during the phantom experiments. Red (numbers 1-2) marks the locations where the internal pressure was measured, green (number 3) marks where the Doppler measurements were performed, and purple (numbers 4-8) indicates where the MRI measurement planes were positioned. Here, the abbreviations AV and MV represent the aortic valve and the mitral valve.

### Magnetic resonance imaging

The mass flow and bulk velocity were measured at the apex, the aortic valve, and in the atrium (Fig. 3) with 2D phase-contrast (PC) MRI at selected measurement planes and analyzed (Segment v3.3 R10057, https://medviso.com/segment/) [20] on a 1.5 T MR clinical scanner (MAGNETOM Sola, Siemens Healthcare, Erlangen, Germany). With 4D (3D + time) PC-MRI the mass flow and velocity were also recorded in the entire domain and analyzed (CAAS MR Solutions 5.2, Pie Medical Imaging BV, Maastricht, The Netherlands) to measure the flow through the same cross-sectional planes as with the 2D PC-MR for comparison. The bulk velocity through the mitral valve (Fig. 3) was also quantified as well as the velocity profile in the ventricle. Additionally, the flow through the phantom was visualized with streamlines based on the 4D PC-MRI data. The valve dynamics were imaged with cine MRI on which the valve opening, defined as the distance between the leaflet tips, was measured during peak flow (Appendix C, Fig. A.2). All measurements were conducted by two observers, repeated 10 times, and averaged. The MRI sequence parameters employed can be found in Appendix B, Table A.1, and first-order background phase correction was performed for all PC-MRI images [21], [22].

### Echocardiography

Commercially available ultrasound systems EPIQ CVx (Philips Medical Systems, Andover, MA, USA) and Vivid E95 (GE Healthcare; Vingmed Ultrasound, Horten, Norway) were used for both 2D and 3D imaging. The velocity through the valve was quantified with continuous wave Doppler Echo on the equivalent to a 3-chamber-view (IntelliSpace Cardiovascular 5, release 5.2, Philips Medical Systems, Best, The Netherlands). Both Echo systems showed equal image quality for velocity time integral (VTI) measurements. However, in the cases with the lowest CO, the spectral signal from the GE system was too weak, thus data from the Philips system was used. The GE system data was averaged over three pump cycles, while the Philips system data was averaged over two pump cycles. The temporal mean and maximum velocities were determined with the VTI based on the spectral velocity curve. The corresponding mean and maximum pressure difference (referred to as the pressure gradient in the clinical setting) was manually calculated with the Bernoulli equation with the density for water (998 kg/m^3^). The Echo systems were not used to determine the pressure difference as these assume the density of blood.

The valve opening, defined as the distance between the leaflet tips, was measured on 2D 3-chamber-view Echo images from the GE system (Appendix C, Fig. A.3). The opening defined as an area was measured on 3D images from the Philips system (4D MV assessment, TOMTEC Imaging Systems GmbH, Unterschleissheim, Germany). For the case with the medium stiff valve and the lowest CO, data from the GE machine was used. All manual Echo measurements were conducted 10 times and averaged, performed by two observers.

### Investigated flow cases

Six different *in vitro* tests were investigated (Table 1). The cases are named accordingly: “Soft”, “Medium”, and “Hard” refer to the valve stiffness and 2.9, 4.4, and 5.4 refer to the CO in l/min.

**Table 1.** Compilation of all flow cases tested in the in vitro experiments where different flow rates are combined with different valve stiffnesses. The cardiac output is abbreviated as CO.

| Case | CO (l/min) | Stiffness (MPa) |
| --- | --- | --- |
| Soft-2.9 | 2.9 | 0.91 |
| Medium-2.9 | 2.9 | 1.89 |
| Soft-4.4 | 4.4 | 0.91 |
| Medium-4.4 | 4.4 | 1.89 |
| Hard-4.4 | 4.4 | 5.13 |
| Medium-5.4 | 5.4 | 1.89 |

For each flow case, all measurements described in the previous section were conducted and are freely available at the Zenodo repository with the assigned digital object identifier DOI: 10.5281/zenodo.10117609 [18]. The STL and STEP files of the phantom geometry, used for 3D printing, are also available [18].

## RESULTS

For visual assessment, snapshots from full-cycle videos obtained from cine MRI, 2D and 3D Echo are depicted to visualize the valve dynamics (Fig. 4).

**Fig. 4.**
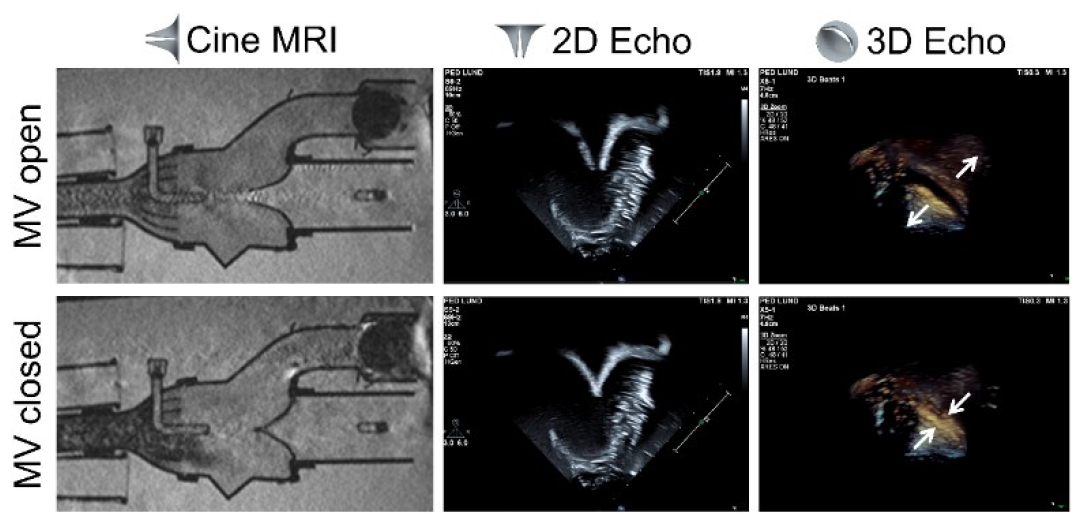
The maximum mitral valve (MV) opening and complete closure imaged with cine MRI, 2D, and 3D Echo, is shown for case Hard-4.4 to exemplify the results. The white arrows mark the leaflet opening and closure in the 3D Echo images and the valve sketches represent the orientation of the valve for each modality.

### Validation of experimental setup

Technical validation of the experimental setup was performed to ensure reliable results. The 2D PC-MRI measurements were conducted at the aortic valve and the atrium of the phantom at the start and the end of each MRI exam of each flow case. A comparison of these two measurements showed a difference of 0.45 ± 0.35 l/min (3.5 %) and 0.86 ± 0.67 l/min (6.7 %) at the aortic valve and atrium, respectively. Pressure measurements in the ventricle were conducted over 17 consecutive pump cycles. A comparison showed a cycle-to-cycle variation of 0.5 % for all cases, except for case Soft-2.9 which exhibited a variation of 4.3 % variation.

### Catheter pressure measurements

The pressure measurements showed similar flow behavior in all six cases. Therefore, only case Medium-4.4 is shown, to exemplify the results (Fig. 5). For the ventricular pressure, differences in amplitude were noted, where a higher peak pressure was observed for higher CO. The atrium was open to the surrounding water reservoir, thus nearly zero (atmospheric) pressure was recorded. The raw data, for all flow cases, is available in the data repository [18].

**Fig. 5.**
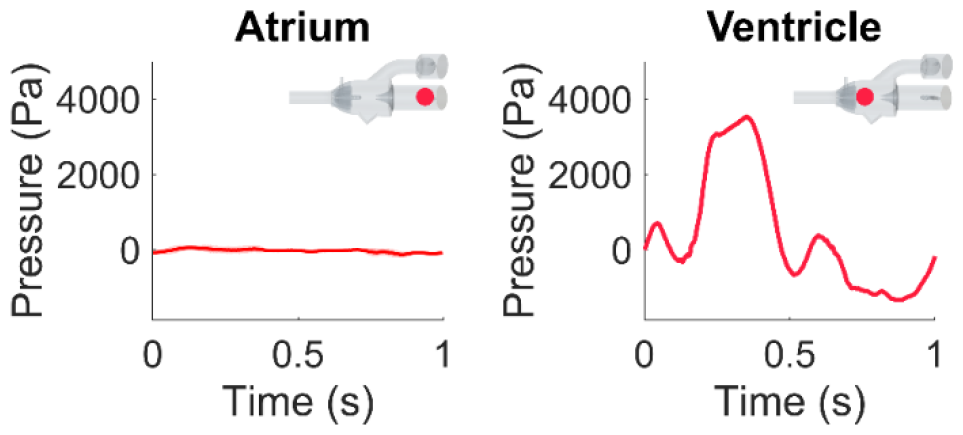
The atrial and ventricular pressure (as indicated by the phantom sketch) in the phantom measured invasively for case Medium-4.4. The data is averaged over 17 pump cycles. Here, the red line represents the average pressure, and the pink background the standard deviation. The sketch of the phantom shows where the pressure was measured.

### MRI measurements

Velocity measurements at the apex, atrium, and aortic valve showed similar behavioral results for all six flow cases both for 2D and 4D PC-MRI measurements. A higher peak velocity was observed for the cases with higher CO; to exemplify the results, case Medium-4.4 is shown (Fig. 6). The remaining measurements are reported in the repository [18]. The flow velocity through the mitral valve increased with increasing CO, measured with 4D PC-MRI data (Fig. 6). A higher peak flow was also observed for increased valve stiffness due to a narrower opening which accelerated the flow.

**Fig. 6.**
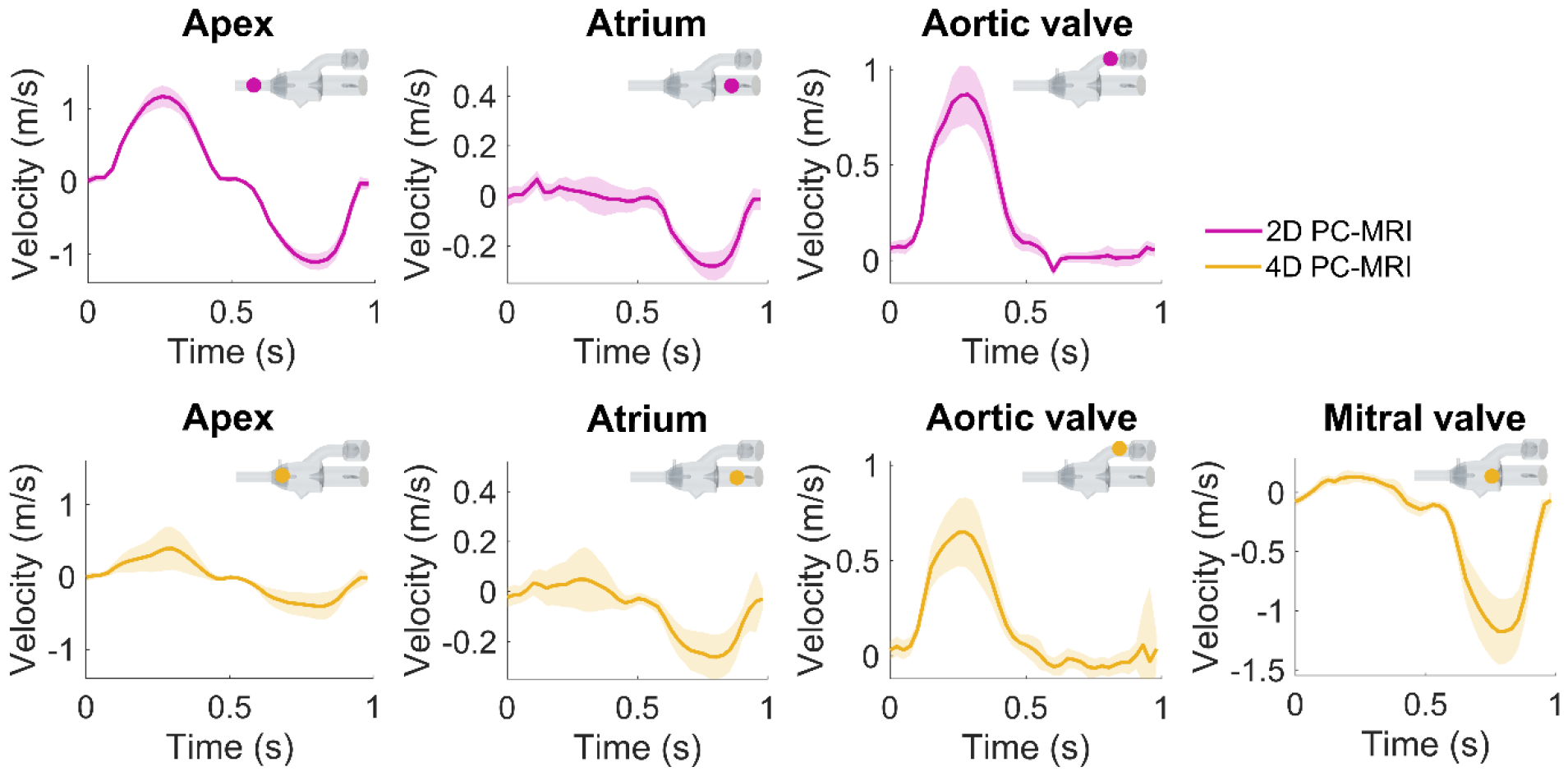
The velocity through cross-sectional planes located at the apex, atrium, aortic valve, and mitral valve, measured with both 2D PC-MRI and 4D PC-MRI, shown for case Medium-4.4. The measurements were spatially averaged across the region of interest, thus the average velocity (dark pink and yellow) is reported together with the standard deviation (light pink and yellow shade) of the measurements. The sketches of the phantom show where the individual measurements were taken, note that the position of the apex measurements differs between the 2D and 4D PC-MRI planes.

In the 4D PC-MRI data, the field of view did not cover the entire phantom, leaving the end of the apex outside the measurable domain. Therefore, the circular cross-sectional plane for flow measurements at the apex was positioned across the baffles of the phantom instead. This resulted in a larger cross-sectional area for the 4D PC-MRI measurement plane compared to the corresponding 2D PC-MRI plane, thus, lower velocities were observed in the 4D PC-MRI apex measurements.

In case Soft-4.4, the softest valve could not withstand the high pressure at CO = 4.4 l/min, which caused it to prolapse. To not damage the silicone valve, the 4D PC-MRI measurements were not conducted for this case.

Velocity profiles were obtained from 4D PC-MRI data along the central lines in y- and z-direction in a cross-sectional plane of the ventricle (Fig. 7c). Figure 7a, d-e shows the data for case Medium-4.4, while other cases are available in the data repository [18]. With the mitral valve closed, a low and uniform velocity profile is observed. However, in the latter half of the pump cycle, a distinct jet through the mitral valve is observed, resulting in recirculation areas. In the velocity profile along the y-direction, a slight dent is present at the center, attributed to the ventricular pressure probe located 3.6 mm downstream. The velocity amplitude peaks at 0.75 s, closely correlated with the pulsatile inlet condition.

**Fig. 7.**
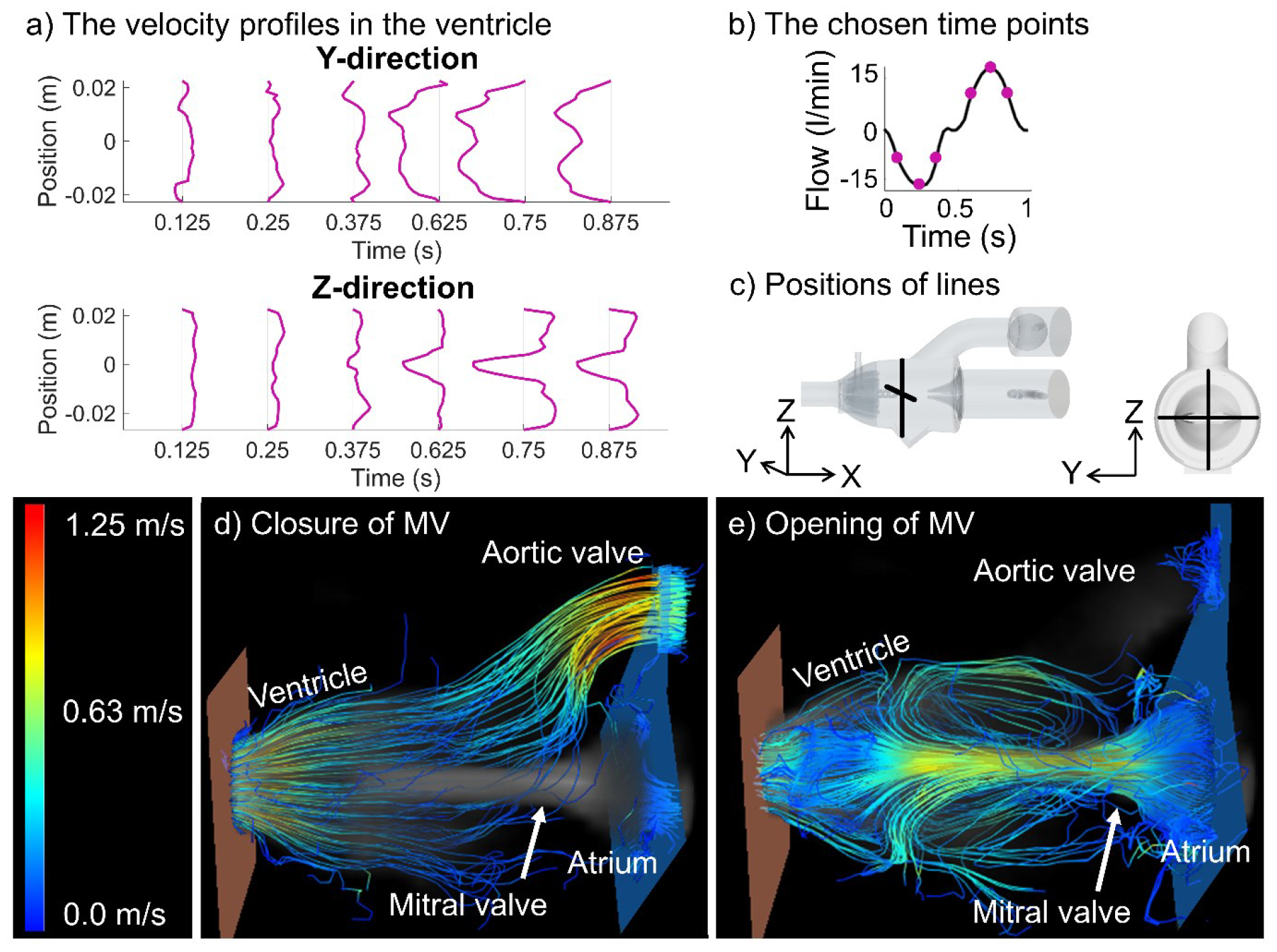
Visual assessment of the flow in the domain, for case Medium-4.4. a) The velocity profile along lines in the y- and z-direction in the ventricle. The y-axes in the plots indicate the position along the measurement lines. b) The graph indicates where in the pump cycle the time points are chosen and (c) shows the position of the measurement within the phantom. The streamlines are shown to visualize the flow inside the phantom for (d) the closure (at t = 0.25 s) and (e) the peak opening (at t = 0.75 s) of the mitral valve (MV).

Streamlines generated from 4D PC-MRI data (Fig. 7d-e) visualize the flow behavior. With the closed mitral valve, water exits through the aortic valve, accelerating through the ventricular outflow tract. At the narrowest part of the open mitral valve, the highest velocity is observed, later decelerated by the baffles at the apex. Recirculation zones within the ventricle are visible, as indicated by the velocity profile.

The valve opening distance was measured on cine MRI data (Appendix C, Fig. A.2). Generally, a larger opening was seen for increased CO and decreased valve stiffness, however, this trend was not seen for cases Soft-2.9 and Medium-2.9. The valve opening measurements varied between 5 mm and 6.1 mm with a standard deviation corresponding to 2.8-5.1 % (Table. 2). The cine MRI measurements showed an interobserver variability of 6.9 ± 4.9 %.

**Table 2.** The valve opening, defined as a distance, measured on cine MRI images. The values are reported as the average value ± the standard deviation of the manual measurements.

| Case | Opening (mm) |
| --- | --- |
| Soft-2.9 | $5.0 \pm 0.22$ |
| Medium-2.9 | $5.7 \pm 0.14$ |
| Soft-4.4 | $6.1 \pm 0.22$ |
| Medium-4.4 | $6.1 \pm 0.31$ |
| Hard-4.4 | $5.8 \pm 0.27$ |
| Medium-5.4 | $6.0 \pm 0.18$ |

### Echo measurements

Doppler Echo velocity measurements through the mitral valve show increased peak velocity for increased CO and stiffness of the valve, due to a smaller opening orifice (Fig. 8). The maximum and mean velocity and pressure difference through the mitral valve were determined from the VTI and the Bernoulli equation (Table 3). The Echo measurements showed an interobserver variability of 24 ± 13 %.

**Table 3.** The mean and maximum velocity and pressure difference (PD) across the mitral valve, measured by Doppler Echo. The values are reported as the average value ± the standard deviation of the manual measurements.

| Case | Mean vel. (m/s) | Max vel. (m/s) | Mean PD (Pa) | Max PD (Pa) |
| --- | --- | --- | --- | --- |
| Soft-2.9 | $0.66 \pm 0.062$ | $1.18 \pm 0.049$ | $217 \pm 1.92$ | $738 \pm 1.20$ |
| Medium-2.9 | $0.76 \pm 0.029$ | $1.43 \pm 0.005$ | $288 \pm 0.42$ | $1090 \pm 0.013$ |
| Soft-4.4 | $1.09 \pm 0.100$ | $1.76 \pm 0.029$ | $593 \pm 4.99$ | $1650 \pm 0.42$ |
| Medium-4.4 | $1.05 \pm 0.010$ | $1.74 \pm 0.031$ | $550 \pm 0.80$ | $1610 \pm 0.48$ |
| Hard-4.4 | $1.22 \pm 0.058$ | $2.10 \pm 0.016$ | $742 \pm 1.68$ | $2360 \pm 0.13$ |
| Medium-5.4 | $1.14 \pm 0.067$ | $1.96 \pm 0.023$ | $648 \pm 2.24$ | $2060 \pm 0.26$ |

**Fig. 8.**
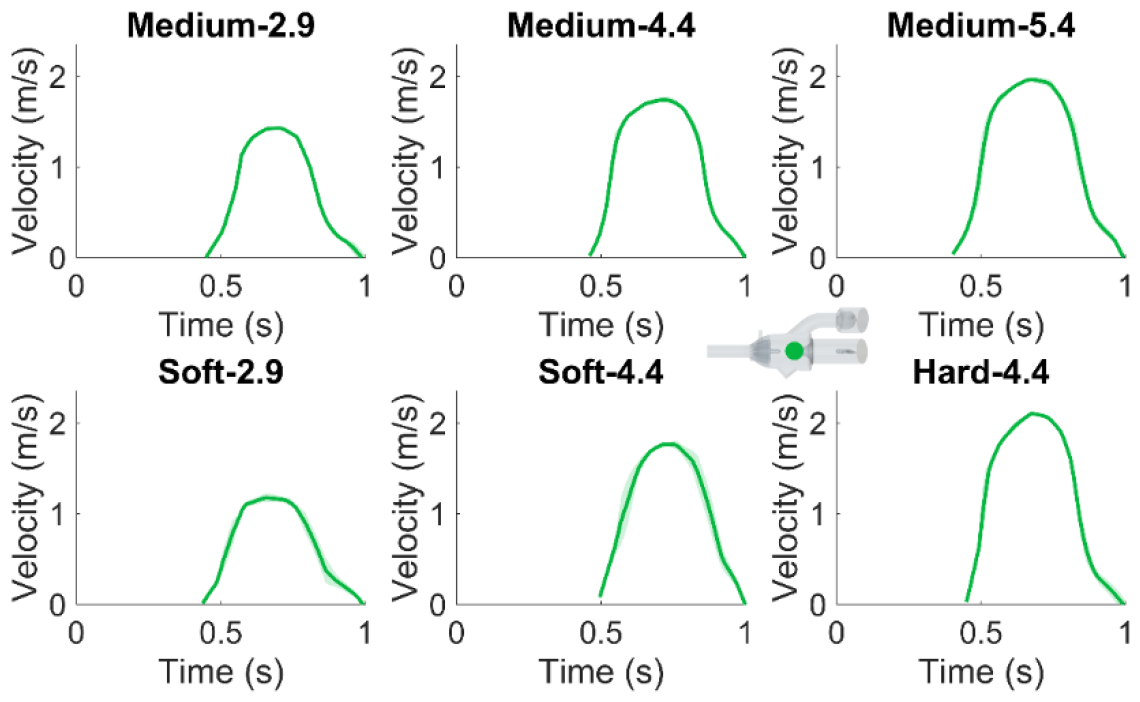
The velocity through the mitral valve over time. The dark green line indicates the average velocity over several pump cycles and the lighter green represents the standard deviation. The sketch of the phantom indicates where the measurements were taken.

For the valve opening distance measurements, ranging from 5.5-6.3 mm with an internal deviation of 2.4-5.1% (Table. 4), an increased opening is observed for increased CO. Similarly, the valve opening area measurements of 0.52-0.69 cm^2^ with a variation of 7.9-14 % (Table. 4), increase for increased CO. Illustrations of how the measurements were conducted are reported in Appendix C, Fig. A.3.

**Table 4.** Manual measurements of the valve opening defined as a distance and as an area, measured 2D and 3D Echo imaging data. All measurements are reported as the average value ± the standard deviation of the manual measurements.

| Case | Opening (mm) | Opening (cm<sup>2</sup>) |
| --- | --- | --- |
| Soft-2.9 | $5.5 \pm 0.27$ | $0.55 \pm 0.05$ |
| Medium-2.9 | $5.6 \pm 0.51$ | $0.52 \pm 0.03$ |
| Soft-4.4 | $6.2 \pm 0.30$ | $0.63 \pm 0.05$ |
| Medium-4.4 | $5.9 \pm 0.30$ | $0.63 \pm 0.05$ |
| Hard-4.4 | $6.3 \pm 0.20$ | $0.64 \pm 0.09$ |
| Medium-5.4 | $6.2 \pm 0.15$ | $0.69 \pm 0.06$ |

## DISCUSSION

This study aimed to create a set of validation data for the development of FSI models, with a phantom setup mimicking the left heart. Pressure measurements, MRI flow measurements, and Echo imaging were used to quantify clinically important measurements 10 typically used for diagnosis of heart valve disease. Specifically, the ventricular and atrial pressure, the velocity within the domain, and the mitral valve opening were quantified in this multi-modal study and are openly provided for the research community [18].

### Validation of experimental setup

A comparison of flow measurements at the start and end of each MRI exam exhibited a low cycle-to-cycle variation, which demonstrated that the setup was reliable over time, with a steady pulsatile flow. This agrees with preliminary experiments from another study on the same motor- and pump setup, which showed a Bland-Altman bias of 1.8 % and an overall MRI error of less than 5 % compared to timer and beaker [24]. The cycle-to-cycle variation in the pressure measurements was smaller than that measured with 2D PC-MRI, which may be attributed to the longer measurement duration for MRI. The pressure was measured for 20 seconds, while the time between 2D PC-MRI scans at the start and end of the experiments was about 30-45 minutes.

### Catheter pressure measurements

The transcatheter approach is considered the gold standard for cardiac pressure measurement [25], [26]. However, catheter placement variability is a common source of error [26]. In our setup, consistent catheter position was secured through fixed attachment points incorporated in the 3D-printed phantom across all measured flow cases. To achieve an *in vitro* test setup, with no malfunctioning valve can be challenging [27]. Previous studies show the challenge in prevention of valve leakage and ensuring reliable pressure measurements [10]. When we calculated the pressure difference across the valve, by subtracting the atrial pressure from the ventricular pressure, the results were comparable with what is reported in literature for valves of different stiffnesses and flow velocities [28].

### MRI measurements

We found discrepancies in the amplitude between 2D and 4D PC-MRI flow measurements. For the apex, this discrepancy is mainly attributed to differences in the position of measurement planes within the phantom, which resulted in different cross-sectional areas and thus deviating velocity readings. However, despite equal positions of the measurement planes at the atrium and aortic valve, differences in velocity measurements were still observed. It has been shown that 4D PC-MRI data may underestimate peak velocities [29], [30], which may account for the observed differences. Additionally, the spatial standard deviation in 4D PC-MRI measurements is larger than in the 2D PC-MRI data, thus a higher variation in velocity in the 4D PC-MRI data was obtained. The lower contrast in 4D PC-MRI data compared to 2D PC-MRI data made accurate delineation of the region of interest for velocity measurements challenging. As the delineated area might include velocity measurements outside the actual flow domain, the average velocity decreases, resulting in an underestimation.

The 4D PC-MRI data allowed qualitative assessment of the ventricular flow during a typical pump cycle. However, the 1.5 mm voxel size in the 4D PC-MRI data could potentially provide insufficient resolution of the velocity jet, thus under-sampling the maximum velocity in the velocity profile, as reported in literature [19]. An alternative is to use particle imaging velocimetry (PIV) for quantification of the velocity profile. This offers higher spatial resolution [10] but requires a transparent flow domain and the material refractive index needs to be accounted for [31]. Despite the spatial resolution limitations, the velocity profile exhibited an expected behavior. Thus, the 4D PC-MRI data can provide a qualitative assessment of the flow behavior, which has the potential to support the development of future FSI models [32].

### Echo measurements

The overall trend and maximum amplitude measured with Doppler align with MRI flow measurements. A higher velocity through the valve is observed for increased CO and valve stiffness, an effect also seen for the mean and maximum pressure differences, consistent with the Bernoulli equation. However, a comparison of catheter pressure measurements and Doppler measurements shows Doppler tends to overestimate the maximum pressure difference across the valve, consistent with literature [33]– [35]. Despite this, Echo measurements are commonly used in the clinical setting, since it is a non-invasive and easily accessible method [36], which further highlights the importance of the conducted measurements in our study as they provide additional insights into Echo measurements.

Identification of the valve was generally easy in most acquired images, despite the potential challenges of imaging 3D-printed materials [37]. However, noisy images, particularly in case Hard-4.4, posed difficulties in accurately identifying valve leaflets and thus measuring distances, which is also reflected in the interobserver variability. Geometric measurements in Echo images are influenced by the angle of the probe relative to the object of interest. Angled cross-sectional planes risk skewed distances and inaccurate geometric measurements. Conversely, 3D volume measurements allow manual positioning of the measurement plane, to ensure proper alignment and avoid skewed distances [38]. In our data, the alignment of the measurement plane was not always perpendicular to the Echo beam itself, which resulted in varying lateral resolution and thus introduced uncertainty in area measurements despite proper alignment in relation to the valve. Nonetheless, the measured opening area follows the same trend as the distance measurements, with larger openings for higher CO and smaller openings for higher valve stiffness.

### Limitations

A limiting factor in this study is that prolapse of the valve was observed in the case Soft-2.9. The soft valve could not withstand the peak pressure that arose at CO 4.4 l/min. Thus, we do not recommend this case to be used for the validation of FSI simulations.

### Conclusions

In this study, a phantom setup mimicking mitral valve dynamics in physiologically inspired conditions, with low cycle-to-cycle variability was built for the validation of cardiac-motivated FSI models. In the *in vitro* experiments parameters relevant to the diagnosis of heart valve disease were measured and quantified by modalities considered the gold standard in clinical practice. Catheter measurements were employed to record ventricular and atrial pressure. The velocity within the phantom was quantified with 2D and 4D (3D + time) PC-MRI measurements, while the valve opening was assessed on cine MRI images as well as 2D and 3D Echo images. Doppler Echo analysis was utilized to quantify the velocity through the mitral valve and the pressure difference across the valve. All obtained results, including the phantom model, have been openly published and made publicly available [18] to facilitate the validation and development of future FSI models for clinical cardiac applications.

## FUNDING AND CONFLICTS OF INTERESTS

The study was supported by grants from the Swedish Heart-Lung Foundation (grant no. 20220737) (PL), the Skåne University Hospital’s Innovation Award (PL), and the Swedish state under the agreement between the Swedish government and the county councils, the ALF-agreement (grant no. 2022-0288) (PL). Additionally, this study was funded by the Swedish Pediatric Heart Foundation and (Hjärtebarnsfonden) (NH) and Skåne University Hospital grant 2022/2023 (NH).

### Declarations

All authors certify that they are not involved in any organization or entity with any financial or non-financial interests that are directly or indirectly related to the work discussed in this manuscript.

## Acknowledgments

The authors would like to thank Peter Paulander for the assistance during the build of the pump setup, Erik Ekbom, and Muris Imsirovic at the 3D center at Skåne University Hospital in Lund for 3D-printing the phantom, and Isabella Silva Barreto for the help with tensile tests. Further, we would like to acknowledge Jonathan Berg for his help with the pressure measurement equipment, and Axel Svenningsson and Pia Sjöberg for their guidance on the software CAAS MR Solutions. Finally, Pie Medical is acknowledged for the CAAS 4D flow analysis research tool.

## Author contributions

The conceptualization of the project was carried out by Nina Hakacova, Petru Liuba, Petter Frieberg, and Lea Christierson. Investigation and data analysis were performed by Lea Christierson, Petter Frieberg, Tania Lala, Nina Hakacova, Johannes Töger, Johan Revstedt, and Hanna Isaksson. Grants from Nina Hakacova and Petru Liuba funded the study. The methodology was developed by Lea Christierson, Petter Frieberg, and Nina Hakacova, who designed the research protocols. Resources were provided by Nina Hakacova, Petter Frieberg, Johannes Töger, and Hanna Isaksson and supervision of the project was provided by Nina Hakacova, Petru Liuba, Johan Revstedt, and Hanna Isaksson. The visualization of data and drafting of the original manuscript were performed by Lea Christierson. The subsequent review, editing, and approval of the final version of the manuscript involved the contributions of all authors.

## APPENDIX

### A. The tensile tests, setup

To quantify the material properties of the silicone material for future material modeling purposes, material tests were performed on N = 6 dog bone samples of stiffness Shore A25 and A40, and N = 5 samples of Shore A60. Uniaxial tensile tests with a preload of 1 N and a loading rate of 6 mm/s, corresponding to a 10 %/s strain rate, were performed until a total of 50 mm displacement was reached. The thickness of the dog bones was the same as the thickness of the mitral valves and the loading rate corresponded to the average loading rate of the valve in the phantom. The dimensions of the dog bone are reported in Fig. A.1a.

#### Results

The tensile tests of the dog bone samples show a bilinear behavior of the silicone material for all three stiffnesses, with the transition region occurring around strains of 0.1-0.3. For strains lower than 0.1, Young’s moduli were 0.91 MPa, 1.89 MPa, and 5.13 MPa for the three different materials. Further, for strains greater than 0.3 Young’s moduli corresponded to 0.51 MPa, 1.25 MPa, and 5.26 MPa (Fig. A.1b).

### B. MRI parameters used for imaging

The phantom was analyzed using cine MRI and 2D and 4D PC-MRI. An overview of all MRI sequence parameters is provided in Table A.1.

**Fig. A.1.**
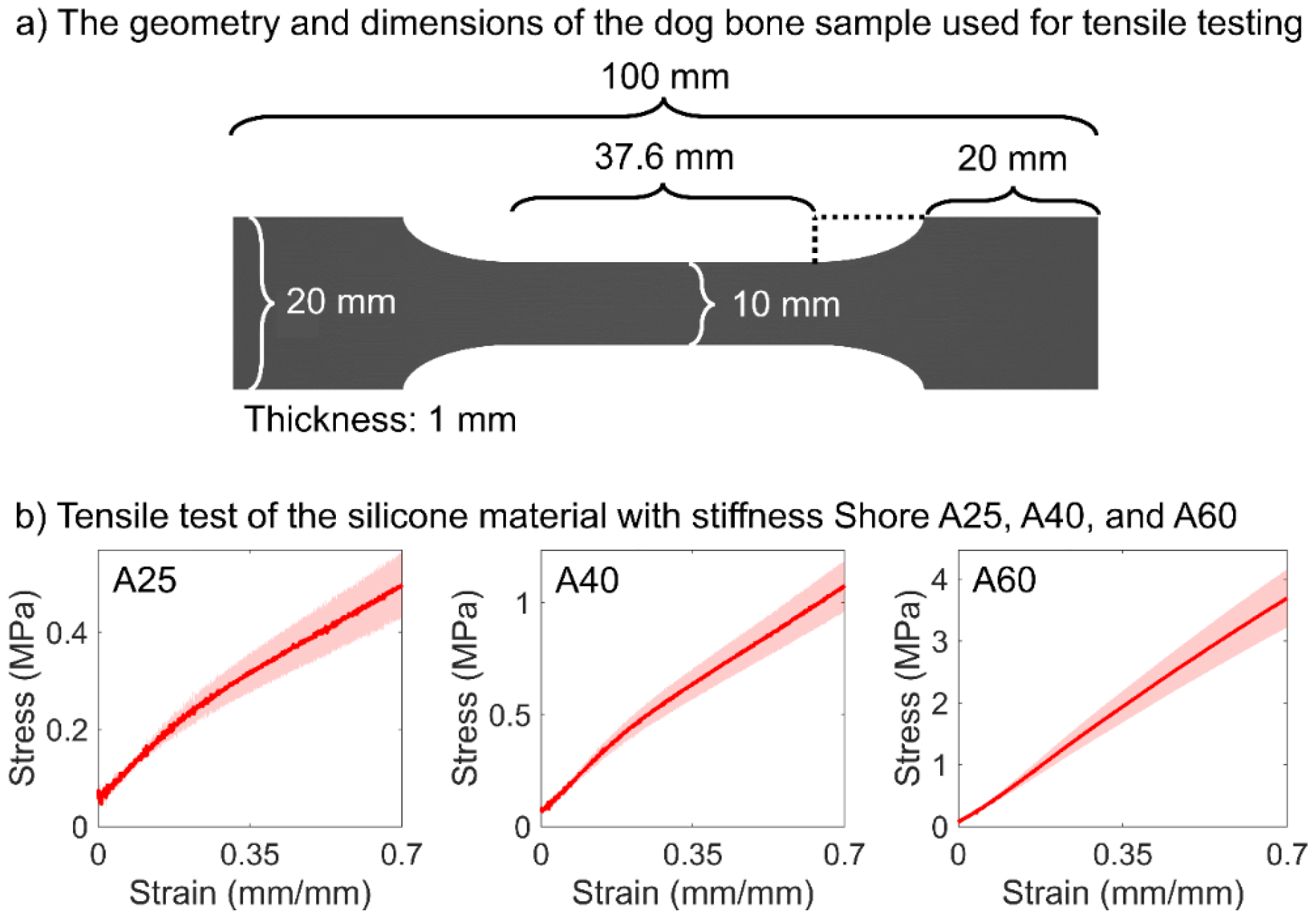
a) The dimensions of the dog bone samples used for material testing. The shape of the dog bone is symmetric about its two in-plane center axes. b) Uniaxial tensile test performed on dog bones of stiffness Shore A25, A40, and A60, respectively. The data is averaged over the number of samples with the mean value shown as a red line and the standard deviation plotted as a pink shadow.

**Table A_1.**
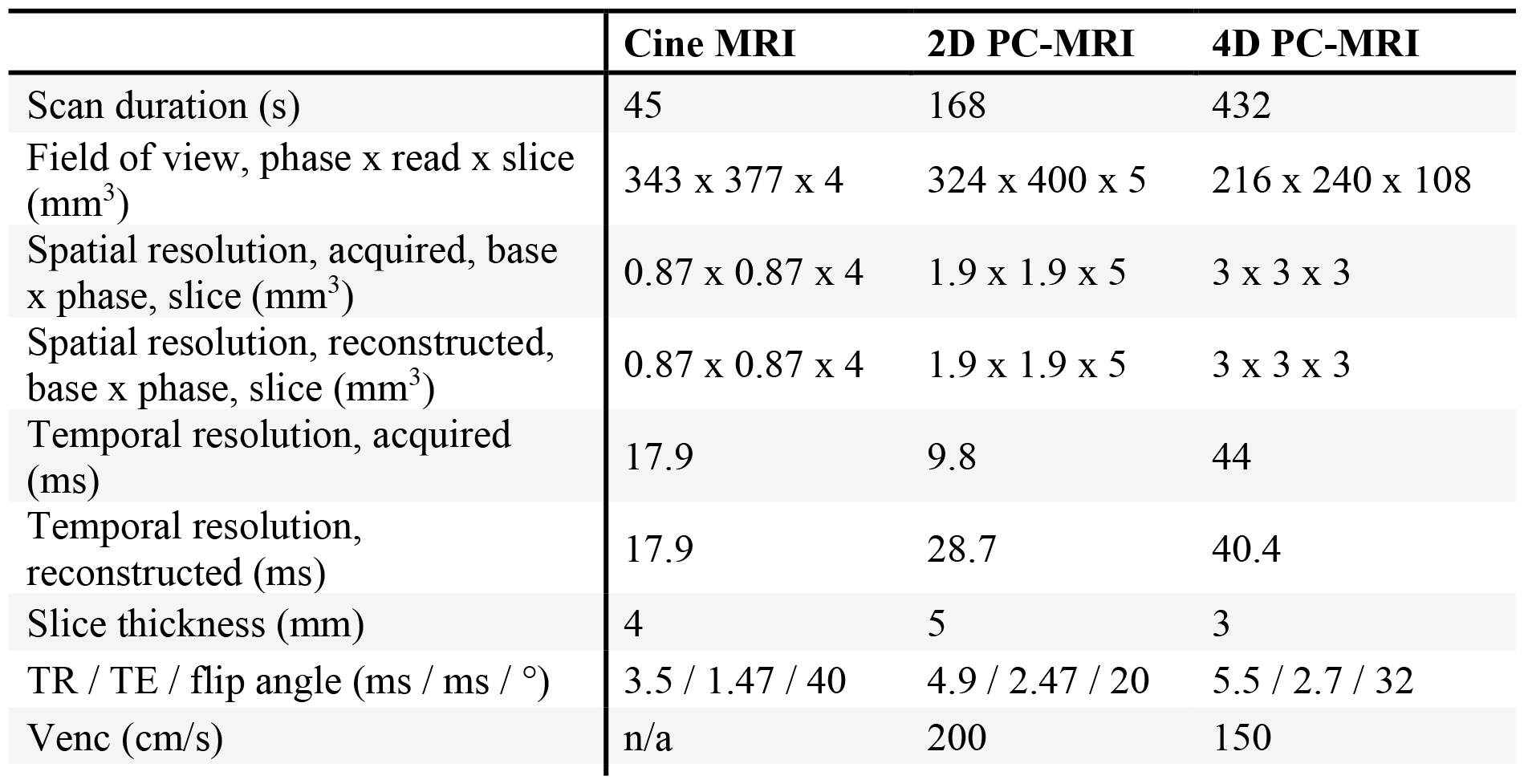
An overview of all MRI sequence parameters.

### C. Measuring the opening of the valve

The valve opening was quantified for all six flow cases based on measurements on cine MRI images (Fig. A.2) and 2D and 3D Echo images (Fig. A.3).

**Fig. A.2.**
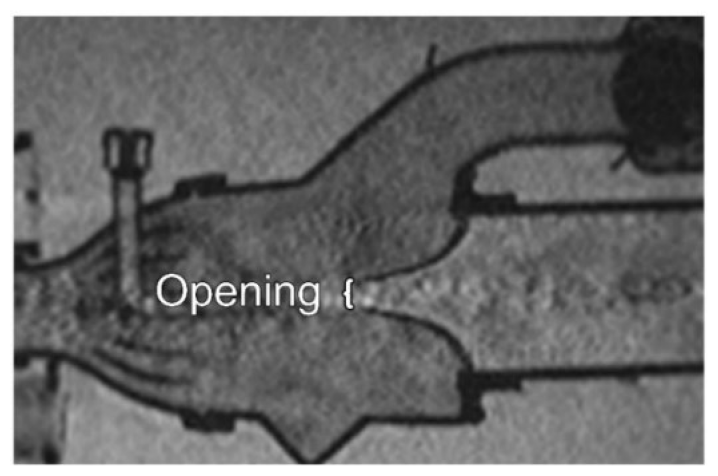
The valve opening, defined as a distance, measured on cine MRI images. The cine MRI image demonstrates where and how the distance measurements were conducted.

**Fig. A_3.**
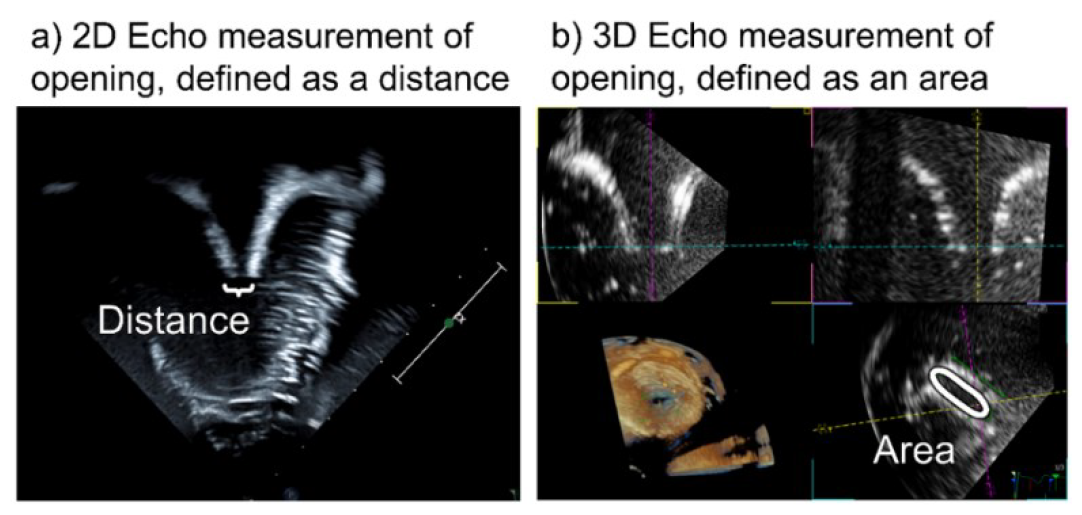
Demonstration of where and how the manual measurements of the valve opening (a) defined as a distance and (b) defined as an area measured on 2D and 3D Echo data were performed.

